# A high-fidelity CRISPR-Cas13 system improves abnormalities associated with C9ORF72-linked ALS/FTD

**DOI:** 10.1101/2023.12.12.571328

**Authors:** Tristan X. McCallister, Colin K. W. Lim, William M. Terpstra, M. Alejandra Zeballos C, Sijia Zhang, Jackson E. Powell, Thomas Gaj

## Abstract

An abnormal expansion of a GGGGCC hexanucleotide repeat in the C9ORF72 gene is the most common genetic cause of amyotrophic lateral sclerosis (ALS) and frontotemporal dementia (FTD), two debilitating neurodegenerative disorders driven in part by gain-of-function mechanisms involving transcribed forms of the repeat expansion. By utilizing a Cas13 variant with reduced collateral effects, we developed a high-fidelity RNA-targeting CRISPR-based system for C9ORF72-linked ALS/FTD. When delivered to the brain of a transgenic rodent model, this Cas13-based platform effectively curbed the expression of the GGGGCC repeat-containing RNA without affecting normal C9ORF72 levels, which in turn decreased the formation of RNA foci and reversed transcriptional deficits. This high-fidelity Cas13 variant possessed improved transcriptome-wide specificity compared to its native form and mediated efficient targeting in motor neuron-like cells derived from a patient with ALS. Our results lay the foundation for the implementation of RNA-targeting CRISPR technologies for C9ORF72-linked ALS/FTD.

## INTRODUCTION

Amyotrophic lateral sclerosis (ALS) is a rapidly progressive, paralytic and invariably fatal disorder characterized by the selective loss of motor neurons in the brain and spinal cord^1^. It also exists on a continuum of conditions that includes frontotemporal dementia (FTD)^2–4^, a syndrome defined by the progressive impairment of cognitive functions due to the degeneration of the frontal and temporal lobes of the brain^5,6^.

An abnormal expansion of a GGGGCC (G_4_C_2_) hexanucleotide repeat in the first intron of the chromosome 9 open-reading frame 72 (C9ORF72) gene is the most common genetic cause of both ALS and FTD^7,8^. To date, three non-exclusive mechanisms have been proposed to explain the pathogenicity of this repeat expansion^9–24^. These include a loss-of-function of the C9ORF72 protein from impaired transcription of the mutant allele^9–11^ and/or an acquired gain-of-function from the bidirectional transcription of sense^12^ and antisense^13^ transcripts carrying the hexanucleotide repeat expansion, which can accumulate in cells as foci^14^, potentially with critical RNA-binding proteins^15–18^. These repeat-containing transcripts can further serve as templates for the synthesis of one of five dipeptide repeat (DPR) proteins^19–22^, which are non-canonically translated^19^ and believed to exert toxic effects on cells^23,24^.

Given the evidence in support of a role for the hexanucleotide repeat-containing RNAs in C9ORF72-linked ALS and FTD – hereafter referred to as C9-ALS/FTD – strategies for silencing their production hold potential for the disorder^25–30^, as they can inhibit the formation of abnormal RNA foci^25,27^ and DPR protein deposits^31,32^. One emerging technology with the capabilities to enable this is Cas13^33–35^, a class 2 type VI CRISPR effector protein that, when complexed with a CRISPR RNA (crRNA) guide molecule carrying complementarity to a target transcript, can cleave it via its intrinsic ribonuclease (RNase) activity. To date, four distinct Cas13 subtypes have been identified and used for gene silencing in eukaryotic cells^33–36^. Among these is the Cas13d nuclease from *Ruminococcus flavefaciens* XPD3002^33^, known as RfxCas13d or CasRx, a CRISPR effector protein that is compact enough to fit within a single adeno-associated virus (AAV) vector to enable its delivery to the central nervous system^37–40^ and whose modification with nuclear localization signal (NLS) sequences can enable it to target transcripts in the nucleus^33^. Given these properties, we hypothesized that RfxCas13d could be used to silence the hexanucleotide repeat-containing RNA and that its action could influence pathological hallmarks of C9-ALS/FTD.

In the present study, we establish a CRISPR-Cas13-based platform for C9-ALS/FTD. Using a dual-luciferase reporter screen designed to assess target engagement and collateral effects, we identify crRNAs for RfxCas13d that facilitate the efficient targeting of the G_4_C_2_ repeat-containing RNA. When delivered to C9-BACexp mice, which harbor the human C9ORF72 gene with a disease-associated expansion of the hexanucleotide repeat, this RfxCas13d-based platform effectively curbed the expression of the G_4_C_2_ repeat-containing RNA without affecting normal C9ORF72 mRNA levels, an outcome we show led to a reduction in RNA foci composed of a transcribed form of the repeat expansion. Further, we demonstrate that a high-fidelity variant of RfxCas13d, which possesses improved transcriptome-wide specificity compared to the native enzyme, can be integrated into our platform to mediate targeting in both motor neuron-like cells derived from a patient with ALS and in C9-BACexp mice, where we find it could inhibit the formation of RNA foci and reverse transcriptional deficits following its *in vivo* delivery by an AAV vector.

Our results thus altogether illustrate the potential of CRISPR-Cas13 technology for C9-ALS/FTD.

## RESULTS

### Programming RfxCas13d to target C9ORF72

The human C9ORF72 gene consists of 11 exons that are transcribed to three major transcript variants, V1, V2 and V3, with the repeat expansion located in intron 1 between exons 1a and 1b (**Fig. 1a**). These three transcript variants produce two protein isoforms: a 222-residue short isoform, known as C9-S, which is translated from V1, and a 481-residue long isoform, C9-L, translated from V2 and V3 (**Fig. 1a**)^10,41^.

**Fig. 1.**
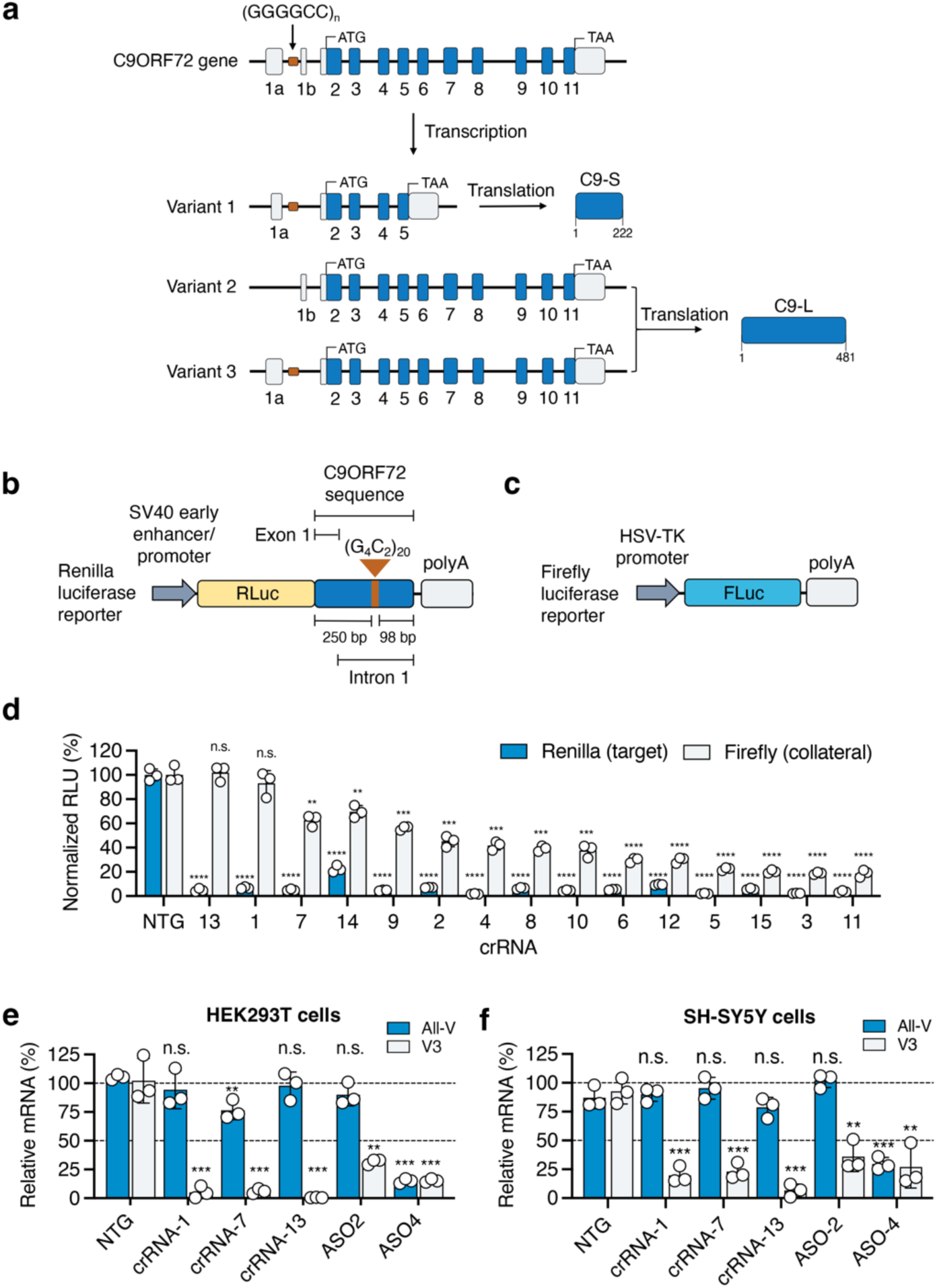
RfxCas13d can be programmed to target C9ORF72. **(a)** Schematic of (top) the C9ORF72 gene and (bottom) the three main mRNA transcript variants expressed from it. V1 produces the short protein isoform (C9-S), while V2 and V3 produce the long protein isoform (C9-L). **(b, c)** Schematic of the dual-reporter system used to evaluate crRNAs. The platform consists of a **(b)** Renilla luciferase-encoding plasmid, pSV40-RLuc, whose 3’ untranslated region (UTR) carries a fragment of the C9ORF72 gene with 20 copies of the hexanucleotide repeat and 250-and 98-base pairs (bps) of the flanking upstream and downstream gene sequences, respectively, and **(c)** a firefly luciferase-encoding plasmid, pHSV-TK-FLuc, which was used as a proxy for collateral cleavage. **(d)** Normalized Renilla and firefly luciferase expression in HEK293T cells transfected with pSV40-RLuc, pHSV-TK-FLuc, and an expression vector encoding RfxCas13d and one of the 15 candidate crRNAs. All values were normalized to cells transfected with pSV40-RLuc, pHSV-TK-FLuc, and an expression vector encoding RfxCas13d with a non-targeted (NTG) crRNA (*n* = 3). **(e, f)** Relative all-V and V3 mRNA in **(e)** HEK293T and **(f)** SH-SY5Y cells transfected with RfxCas13d and crRNAs 13, 7, and 1 or a NTG crRNA or one of two ASOs (*n* = 3). All values from HEK293T and SH-SY5Y cells were normalized to untreated cells. Values indicate means and error bars indicate SD. **P < 0.01, ***P < 0.001, ****P < 0.0001; one-tailed unpaired t-test comparing each crRNA to the NTG crRNA. All data points are biologically independent samples.

Both V1 and V3 encode the repeat expansion, while transcription of V2, the predominant variant thought to account for ∼85-95% of the C9ORF72 transcripts in the brain^25,42^, is initiated downstream of the repeat (**Fig. 1a**)^13^. Because the partial loss of the C9ORF72 protein has been hypothesized as a potential source of the pathogenicity of the repeat expansion^43,44^, we sought to use RfxCas13d to target either exon 1a or intron 1 to preferentially silence V1 and V3 (**Fig. 1a**), a strategy we expected would spare V2, a major source of the C9-L protein.

To facilitate the design of crRNAs to silence the repeat-containing RNA, we utilized the Cas13 Design Resource, an online tool that can predict active crRNAs for RfxCas13d^45,46^. Using this resource, we searched the exon 1a and the intron 1 sequences flanking the hexanucleotide repeat before selecting the 15 highest ranking crRNAs for detailed testing (**Supplementary Fig. 1**). We then created a dual-luciferase reporter screen to determine the ability of the crRNAs to target their respective sequences. This platform consists in part of a Renilla luciferase transgene whose 3’ untranslated region (UTR) is fused to a fragment of the C9ORF72 gene encoding 20 copies of the hexanucleotide repeat with 250-and 98-base pairs (bps) of the flanking upstream and downstream gene sequences, respectively (**Fig. 1b**). When complexed with a functional crRNA, RfxCas13d is expected to target the 3’ UTR of the Renilla luciferase transcript, which in turn is anticipated to decrease its expression.

In addition, because RfxCas13d has the potential to degrade non-specific RNAs via its *trans*-cleavage activity^47,48^, we utilized firefly luciferase as a secondary reporter to measure collateral effects (**Fig. 1c**). Importantly, by this approach, firefly luciferase expression is expected to decrease if *trans*-cleaved by RfxCas13d. As the extent of *trans*-cleavage has been observed to depend in some cases on the crRNA^49,50^, we reasoned this strategy could aid in the identification of crRNA targeting sequences that minimize this effect.

To test the crRNAs, we transfected human embryonic kidney (HEK) 293T cells with each reporter plasmid and an expression vector encoding a RfxCas13d variant with two NLS sequences, one attached to the N-terminus and the other to the C-terminus, and one of the 15 candidate repeat-targeting crRNAs. From this screen, we found that all 15 crRNAs decreased Renilla luciferase by >75% at 72 hr post-transfection (P < 0.0001) and that 14 of the 15 crRNAs decreased it by at least 95% (P < 0.0001; **Fig. 1d**); however, we found that the majority of the crRNAs also triggered a reduction in the expression of firefly luciferase (**Fig. 1d**), indicating that they exerted collateral effects. This notwithstanding, we identified two crRNAs, crRNAs 13 and 1, that decreased Renilla luciferase by ∼95% (P < 0.0001) and had no significant effect on firefly luciferase compared to the controls (P > 0.05; **Fig. 1d**).

We next evaluated if RfxCas13d could target C9ORF72 mRNA in HEK293T cells. More specifically, we tested RfxCas13d with crRNAs 13, 7 and 1 which, among the five crRNAs found to reduce firefly luciferase expression by less than two-fold (**Fig. 1d**), possessed the most favorable targeting scores, a number we defined as the ratio of Renilla (i.e., target) to firefly (i.e., collateral) luciferase expression (**Supplementary Fig. 2**). To determine targeting, we used validated qPCR probes^32^ to measure: (i) V3, which served as our proxy for the G_4_C_2_ repeat-containing RNA, and (ii) the all-variant (all-V) pool, which consists predominately of V2^25,42,51^ and thus is not expected to be perturbed by our approach.

Compared to cells transfected with RfxCas13d and a non-targeted crRNA, we measured by qPCR that each of the three repeat-targeting crRNAs decreased the relative abundance of V3 mRNA by ∼90-95% (P < 0.0001 for all; **Fig. 1e**). Importantly, relative to the same controls, we measured that these crRNAs either had no effect on all-V (crRNAs 13 and 1; P > 0.05) or a limited effect on it (crRNA-7; **Fig. 1e**).

As an additional control, we tested in HEK293T cells two previously validated antisense oligonucleotides (ASOs) for C9ORF72: ASO-2, which binds intron 1 and selectively targets V1 and V3^27^, and ASO-4, which binds exon 2 and thus targets the three main transcript variants^27^. As expected, qPCR revealed that ASO-2 lowered only V3 (P < 0.01) and that ASO-4 decreased both V3 and all-V (P < 0.001 for both; **Fig. 1e**), reinforcing the validity of our measurement methods.

We next evaluated if RfxCas13d could preferentially target V3 in a second human cell line, SH-SY5Y neuroblastoma cells. Compared to cells transfected with RfxCas13d and a non-targeting crRNA, we found by qPCR that crRNAs 13, 7 and 1 decreased V3 mRNA by 80-95% (P < 0.001 for all; **Fig. 1f**) and that each of the three crRNAs had no significant effect on all-V (P < 0.05 for all; **Fig. 1f**). Consistent with our prior control measurements, ASO-2 was again found to only lower V3 in SH-SY5Y cells (P < 0.01), while ASO-4 decreased both V3 and all-V (P < 0.001 for V3 and P < 0.01 for all-V; **Fig. 1f**).

In sum, we find that a dual-luciferase reporter screen designed to measure collateral effects can be used to identify crRNAs for C9ORF72, and that RfxCas13d can preferentially target V3, a C9ORF72 transcript variant that carries the hexanucleotide repeat, in human cells.

### RfxCas13d targeting can decrease the G_4_C_2_ repeat-containing RNA and reduce RNA foci formation in a C9-ALS/FTD mouse model

We next sought to determine whether RfxCas13d could target the hexanucleotide repeat-containing transcript in a mouse model of C9-ALS/FTD, specifically C9-BACexp mice^52^, which harbor a bacterial artificial chromosome (BAC) encoding the full-length human C9ORF72 gene with ∼100-1,000 copies of the G_4_C_2_ repeat. Similar to other rodent models of C9-ALS/FTD^53^, C9-BACexp mice do not manifest an overt motor phenotype, however, they do develop several hallmarks of the disorder, including the accumulation of RNA foci.

To deliver RfxCas13d, we used the AAV vector variant PHP.eB^54^ which, when injected to the brain parenchyma, can transduce cells to a similar extent as AAV9^55^ but, in the experience of our laboratory, packages at a higher titer. The hippocampus (HPC) and motor cortex (MC) of two-month-old C9-BACexp mice were thus injected with 2 x 10^10^ genome copies (GCs) of an AAV-PHP.eB vector carrying a CBh-driven RfxCas13d variant with one of three repeat-targeting crRNAs, crRNA 13, 7 or 1 (i.e., AAV-PHP.eB-RfxCas13d-crRNA-13, -7 or -1), or a non-targeting crRNA (i.e., AAV-PHP.eB-RfxCas13d-NTG; **Fig. 2a**).

**Fig. 2.**
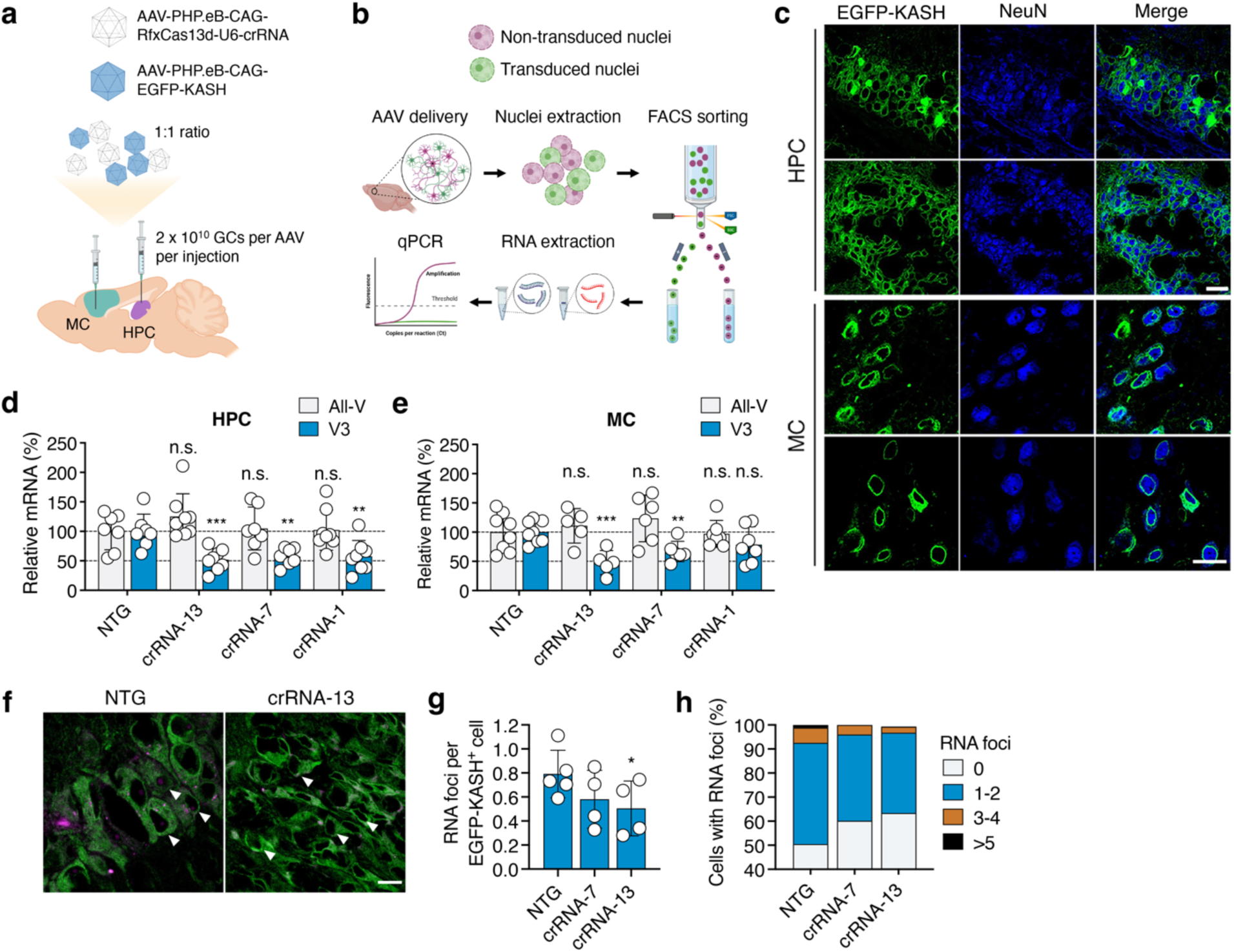
RfxCas13d can target the G_4_C_2_ repeat-containing RNA and reduce the formation of RNA foci in the brain of C9-BACexp mice. **(a)** Cartoon illustrating the injection scheme. **(b)** Overview of the experimental plan to analyze C9ORF72 mRNA in EGFP-KASH^+^ nuclei isolated by fluorescence-activated cell sorting (FACS). **(c**) Representative immunofluorescent staining of the hippocampus (HPC) and motor cortex (MC) in C9-BACexp mice two-months after injection with 2 x 10^10^ GCs each of AAV-PHP.eB-RfxCas13d-crRNA and AAV-PHP.eB-EGFP-KASH. Scale bar, 20 µm. **(d, e)** Relative all-V and V3 mRNA in **(d)** the HPC and **(e)** MC of EGFP-KASH^+^ nuclei from C9-BACexp injected with AAV-PHP.eB-RfxCas13d-crRNA-13, -7, -1, or -NTG with AAV-PHP.eB-EGFP-KASH (*n* ≥ 7). **(f)** Representative RNA FISH to detect the G_4_C_2_ repeat RNA (purple) from the HPC of C9-BACexp mice injected with AAV-PHP.eB-RfxCas13d-crRNA-13 or -NTG and AAV-PHP.eB-EGFP-KASH. Arrowheads indicate representative cells. Scale bar, 15 µm. **(g, h)** Quantification of **(g)** the number of RNA foci per EGFP-KASH^+^ cell in the HPC and **(h)** the percentage of EGFP-KASH^+^ cells with 0, 1-2, 3-4, or >5 foci (*n* ≥ 4). **(g, h)** 58-455 cells were counted per animal. 398, 845, and 467 cells total were counted for AAV-PHP.eB-RfxCas13d-crRNA-7, crRNA-13, and -NTG respectively (*n* ≥ 4). RNA foci measurements were conducted by a blinded investigator. Values indicate means and error bars indicate SD. *P < 0.05, **P < 0.01, ***P < 0.001; one-tailed unpaired t-test comparing each crRNA to the NTG crRNA. All data points are biologically independent samples.

To facilitate the isolation of transduced nuclei for a more detailed analysis of RfxCas13d-mediated outcomes, we co-injected C9-BACexp mice with 2 x 10^10^ GCs of a second AAV-PHP.eB vector encoding a CBh-driven EGFP variant fused to KASH (Klarsicht/ANC-1/Syne-1 homology; AAV-PHP.eB-EGFP-KASH), a domain that promotes EGFP localization to the outer nuclear membrane^56^, which can enable the isolation of transduced nuclei by fluorescence-activated cell sorting (FACS)^57,58^ (**Fig. 2a-b**).

Based on an immunofluorescent analysis conducted at two-months post-injection, we observed strong EGFP-KASH expression at the injection sites of C9-BACexp mice treated with each AAV formulation. Within the HPC and MC, we determined that ∼91% and ∼53% of the cells positive for the pan-neuronal marker NeuN, respectively, were positive for EGFP-KASH (**Fig. 2c** and **Supplementary Fig. 3**) and that ∼72% and ∼75% of the EGFP-KASH^+^ cells in the HPC and MC, respectively, were positive for RfxCas13d by its hemagglutinin (HA) epitope (**Supplementary Fig. 4**). Despite relying on the ubiquitous CBh promoter to drive its expression, EGFP-KASH was largely confined to NeuN^+^ cells, as we observed limited expression in GFAP^+^ astrocytes and Iba1^+^ microglia in both the HPC and MC (**Supplementary Fig. 5**).

We next used FACS to isolate EGFP-KASH^+^ nuclei from the HPC and MC of injected C9-BACexp mice to determine the abundance of V3 and all-V by qPCR (**Fig. 2b**). Relative to EGFP-KASH^+^ cells from mice injected with AAV-PHP.eB-RfxCas13d-NTG, we measured that each repeat-targeting crRNA effectively decreased V3 mRNA (**Fig. 2d-e**), with the most potent and consistent crRNA, the exon 1a-targeting crRNA-13, found to suppress V3 by ∼48% in the HPC (P < 0.001; **Fig. 2d**) and ∼52% in the MC (P < 0.001; **Fig. 2e**). Critically, all three repeat-targeting crRNAs were also found to have no effect on all-V (P > 0.05 for each for the HPC and MC; **Fig. 2d-e**), indicating that RfxCas13d preferentially targeted V3 *in vivo*.

Using fluorescence *in situ* hybridization (FISH), we next determined whether targeting the repeat RNA with RfxCas13d affected RNA foci formation in C9-BACexp mice, specifically in the HPC, where they develop in 40-60% of cells^52^. Given their improved targeting compared to crRNA-1, we quantified RNA foci in EGFP-KASH^+^ cells only for mice treated with RfxCas13d and crRNAs 13 and 7.

Compared to animals infused with control vector, C9-BACexp mice injected with AAV-PHP.eB-RfxCas13d-crRNA-13 and AAV-PHP.eB-RfxCas13d-crRNA-7 had a ∼37% and a ∼31% decrease, respectively, in the mean number of foci per cell positive for the G_4_C_2_ repeat-containing RNA (P < 0.05 for crRNA 13; P = 0.09 for crRNA 7; **Fig. 2f-g**). Additionally, we measured a shift in the distribution of the number of foci per cell for the treated animals, with C9-BACexp mice infused with AAV-PHP.eB-RfxCas13d-crRNA-13 and AAV-PHP.eB-RfxCas13d-crRNA-7 each found to have an 11-13% increase in the percentage of EGFP-KASH^+^ cells with no detectable foci and an approximately two-fold decrease in the percentage of EGFP-KASH^+^ cells with 3-4 foci compared to the controls (**Fig. 2h**).

Thus, we find that RfxCas13d can be delivered to C9-BACexp mice by an AAV vector variant, where it could preferentially suppress the G_4_C_2_ repeat-containing RNA, which affected the formation of RNA foci in neurons.

### High-fidelity RfxCas13d can target C9ORF72 with improved transcriptome-wide specificity

Because of their risk for inducing collateral effects^47,48^, high-fidelity forms of Cas13 that possess a reduced capacity to *trans*-cleave non-target RNAs have been developed^50^. Given their potential for enhancing the specificity of silencing, we determined the ability of these higher fidelity variants to target C9ORF72.

To this end, we transfected HEK293T cells with an expression vector encoding the native RfxCas13d protein or one of two high-fidelity variants^50^, RfxCas13d-N2V7 and RfxCas13d-N2V8 (**Fig. 3a**) with either crRNA-13, the top-performing repeat-targeting crRNA from our first study, or a non-targeted crRNA.

**Fig 3.**
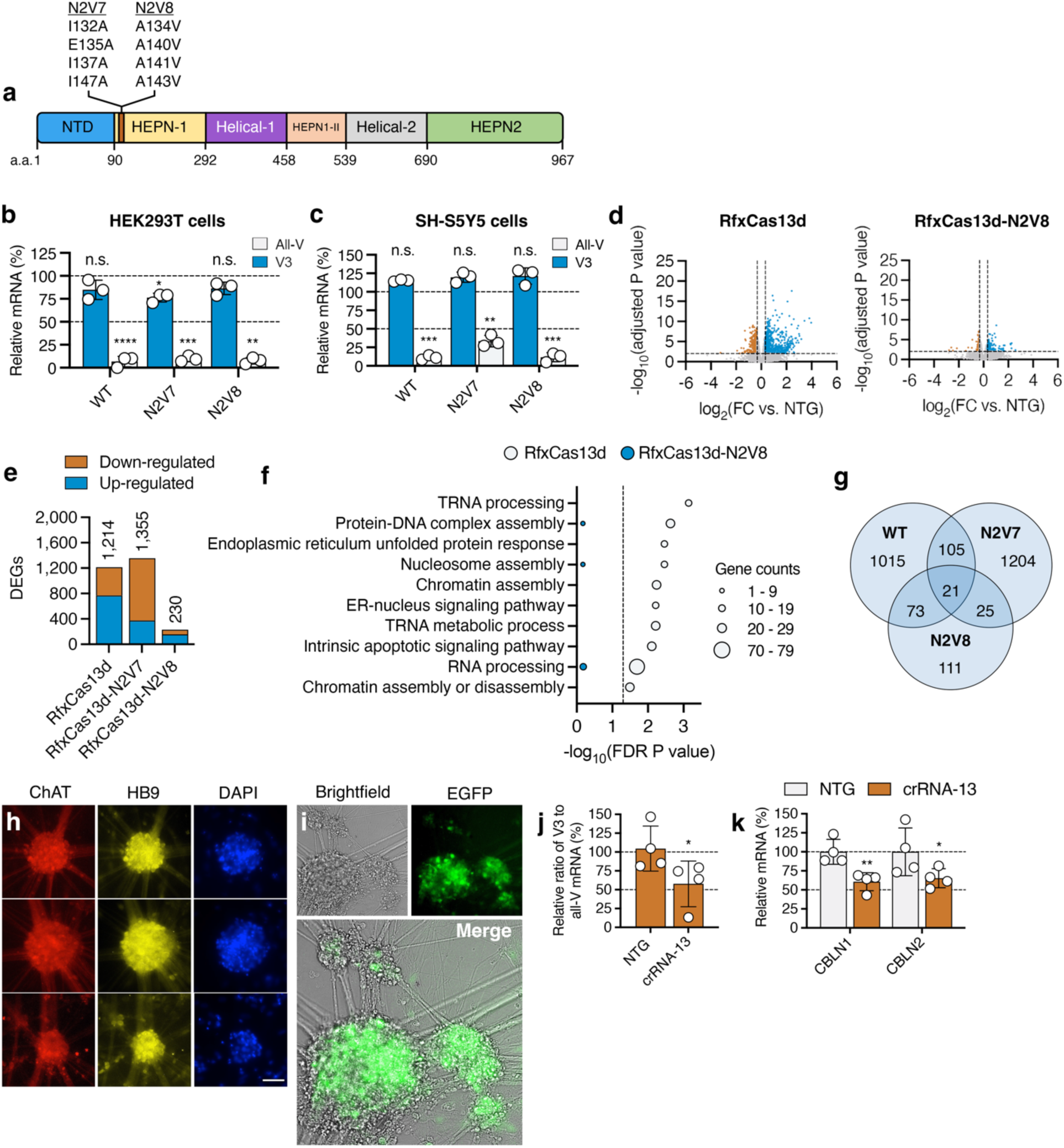
High-fidelity RfxCas13d has improved specificity and can mediate targeting in cells derived from an ALS patient. **(a)** RfxCas13d domain organization and RfxCas13d-N2V7 and RfxCas13d-N2V8 mutations. (b, c) Relative all-V and V3 mRNA in (b) HEK293T and (c) SH-SY5Y cells transfected with RfxCas13d, RfxCas13d-N2V7 and RfxCas13d-N2V8 with crRNA-13 or a non-targeted (NTG) crRNA. All values normalized to untreated cells (*n* = 3). (d) Volcano plot of the RNA-seq analysis comparing HEK293T cells transfected with (left) RfxCas13d or (right) RfxCas13d-N2V8 with crRNA-13 to each variant with NTG crRNA (*n* = 3). Lines denote a >1.25-fold change (FC) and an FDR-adjusted P < 0.01. (e) Number of differentially expressed genes (DEGs) [>1.25-FC, FDR-adjusted P < 0.01] from (d). (f) Gene ontology (GO) and biological process (BP) term analysis for the DEGs in (d). Line denotes FDR-adjusted P < 0.05. (g) Venn diagram of overlapping DEGs. (h) Immunostaining of neurospheres. Cells positive for Hb9 and choline acetyltransferase (ChAT). Scale bar, 75 µm. (i) Brightfield and fluorescent images of neurospheres seven days after treatment with AAV-PHP.eB-EGFP-KASH. (j, k) Relative (j) V3 to all-V mRNA and (k) CBLN1 and CBLN2 mRNA in neurospheres treated with AAV-PHP.eB-RfxCas13d-N2V8-crRNA-13 or -NTG (*n* = 4). Values indicate means and error bars indicate SD. *P < 0.05, **P < 0.01, ***P < 0.001, ****P < 0.0001; (b, c) one-tailed unpaired t-test comparing crRNA-13 to NTG crRNA for each variant; (j, k) one-tailed unpaired t-test comparing crRNA-13 to NTG crRNA (b, c, j, k). All data points are biologically independent samples.

Based on qPCR, both RfxCas13d-N2V7 and -N2V8 decreased V3 mRNA in HEK293T cells (P < 0.001 for RfxCas13d-N2V7 and P *<* 0.01 for -N2V8; **Fig. 3b**), with RfxCas13d-N2V8 found to reduce its expression by >90% compared to cells transfected with a non-targeted crRNA, an effect on par with the native protein (**Fig. 3b**). Similar to the native enzyme, RfxCas13d-N2V8 was also found to have no effect on all-V in HEK293T cells (P > 0.05; **Fig. 3b**).

We next tested RfxCas13d-N2V7 and -N2V8 in SH-SY5Y cells, where we observed by qPCR that both variants effectively decreased V3 mRNA (P < 0.05 for both; **Fig. 3c**) and had no effect on all-V (P > 0.05; **Fig. 3c**), though RfxCas13d-N2V8 was found to target V3 more efficiently (∼89% decrease) than RfxCas13d-N2V7 (∼63% decrease).

To further interrogate the effect of targeting V3 by RfxCas13d, we used western blot to measure C9-L, the full-length C9ORF72 protein isoform, in HEK293T cells. Because our approach preferentially targets V3 and because C9-L is encoded by V2 (**Fig. 1a**), we expected that RfxCas13d and its high-fidelity counterparts would have no effect, or a minimal effect, on the relative abundance of C9-L. Consistent with this reasoning, no significant difference in the C9-L protein was measured in cells transfected with RfxCas13d, RfxCas13d-N2V7, or RfxCas13d-N2V8 with crRNA-13 relative to the respective controls (P > 0.05 for all; **Supplementary Fig. 6**). C9-S, the short C9ORF72 protein isoform encoded by V3, could not be detected, and thus was not analyzed.

We next used RNA-seq to compare the transcriptome-wide specificities of RfxCas13d with its high-fidelity counterparts. When programmed with crRNA-13, the native RfxCas13d protein was found to perturb the expression of 1214 genes in HEK293T cells (>1.25-fold change [FC]; false discovery rate [FDR]-adjusted P < 0.05), while RfxCas13d-N2V8 affected only 230 genes, a ∼5.8-fold decrease (**Fig. 3d-e** and **Supplementary Data 1 and 2**). Interestingly, we found that RfxCas13d-N2V7 influenced a similar number of genes (1355) as the native protein (**Fig. 3e** and **Supplementary Data 1 and 2**), indicating it possessed decreased transcriptome-wide specificity relative to RfxCas13d-N2V8 when paired with crRNA-13.

Through an over-representation analysis of gene ontology (GO) and biological process (BP) terms, we next compared the themes enriched for the genes affected by the native RfxCas13d protein and RfxCas13d-N2V8. For RfxCas13d, we observed enrichment (FDR-adjusted P < 0.05) for ten biological functions (**Fig. 3f** and **Supplementary Data 3**), including RNA processing, protein-DNA complex assembly, intrinsic apoptotic signaling and chromatin assembly and disassembly, several of which have been linked to RfxCas13d and its collateral effects^50^. However, no enrichment (FDR-adjusted P > 0.05) was observed for these or any other themes in cells transfected with RfxCas13d-N2V8 (**Fig. 3f** and **Supplementary Data 4**).

As expected, given our approach spares V2, the predominant C9ORF72 transcript variant, we observed no enrichment for function(s) related to the C9ORF72 protein for any tested RfxCas13d variant. Interestingly, unlike for RfxCas13d-N2V7, the majority (68%) of the genes affected by RfxCas13d-N2V8 were up-regulated (**Fig. 3e**); however, no themes (FDR-adjusted P > 0.05) were identified for these differentially expressed genes (DEGs).

Last for this analysis, we analyzed the shared DEGs between the native RfxCas13d protein, RfxCas13d-N2V8 and RfxCas13d-N2V7. In total, only 21 DEGs were shared among all three variants, though 94 DEGs were shared between RfxCas13d and RfxCas13d-N2V8 (**Fig. 3g** and **Supplementary Data 2**), with a GO and BP term analysis found to reveal an enrichment in functions related to the positive regulation of cellular component organization and epithelial cell differentiation (**Supplementary Data 5)**.

As our RNA-seq study revealed that RfxCas13d-N2V8 possessed improved transcriptome-wide specificity compared to its native form and RfxCas13d-N2V7, we next determined its ability to target V3 in a more physiologically relevant model: iPSC-derived motor neuron-like cells from a 74-year-old female ALS patient with >145 copies of the hexanucleotide repeat in the C9ORF72 gene.

Starting from neural progenitor cells, we conducted a two-week differentiation protocol that led to the formation of HB9^+^ and ChAT^+^ neurospheres (**Fig. 3h**), which we treated with AAV-PHP.eB encoding a CBh-driven RfxCas13d-N2V8 variant with crRNA-13 (i.e., AAV-PHP.eB-RfxCas13d-N2V8-crRNA-13) or a non-targeted crRNA (i.e., AAV-PHP.eB-RfxCas13d-N2V8-NTG) at a multiplicity of infection (MOI) of ∼2 x 10^6^. To visualize transduction, we separately treated HB9^+^ and ChAT^+^ neurospheres with an AAV-PHP.eB encoding a CBh-driven EGFP-KASH at the same MOI.

After determining that EGFP was expressed in cells within the neuron-like clusters (**Fig. 3i**), we used qPCR to measure V3 and all-V from the treated neurospheres. To account for variation in C9ORF72 expression between wells, we determined the relative ratio of V3 to all-V for each sample, finding that neurospheres treated with AAV-PHP.eB-RfxCas13d-N2V8-crRNA-13 had a nearly two-fold decrease in relative V3 mRNA compared to cells treated with AAV-PHP.eB-RfxCas13d-N2V8-NTG (P < 0.05; **Fig, 3j**).

We next asked if RfxCas13d-N2V8 could reverse, or partially reverse, disease-associated transcriptional alterations in neuron-like cells. To answer this, qPCR was used to measure the expression of CBLN1 and CBLN2, two members of the cerebellin family of proteins that normally contribute to the formation and function of synapses but are up-regulated in neuron-like cells derived from patients with C9-ALS^59,60^. Based on qPCR, neurospheres treated with AAV-PHP.eB-RfxCas13d-N2V8-crRNA-13 had a ∼40% (P < 0.001) and a ∼35% (P < 0.05) decrease in CBLN1 and CBLN2 mRNA, respectively, compared to cells treated with the control vector (**Fig. 3k**), indicating that RfxCas13d-N2V8 targeting influenced the expression of genes previously found to be affected in C9-ALS/FTD.

Thus, these results demonstrate that high-fidelity forms of RfxCas13d can be used to target V3 and possess improved transcriptome-wide specificity compared to the native enzyme. We also find that that a high-fidelity RfxCas13d variant can reduce the relative abundance of the G_4_C_2_ repeat-containing RNA in neuron-like cells derived from a C9-ALS patient.

### High-fidelity RfxCas13d can target the G_4_C_2_ repeat RNA and revert transcriptional deficits *in vivo*

Given its improved targeting capabilities compared to the native protein, we next evaluated if RfxCas13d-N2V8 could target the hexanucleotide repeat-containing RNA *in vivo*. Mirroring our earlier study, we injected the HPC and MC of two-month-old C9-BACexp mice with 2 x 10^10^ GCs of AAV-PHP.eB-RfxCas13d-N2V8-crRNA-13 or AAV-PHP.eB-RfxCas13d-N2V8-NTG. Further, to facilitate the isolation of transduced nuclei for a more detailed analysis of RfxCas13d-N2V8-mediated outcomes, each site was co-injected with 2 x 10^10^ GCs of AAV-PHP.eB-EGFP-KASH (**Fig. 4a**).

**Figure 4.**
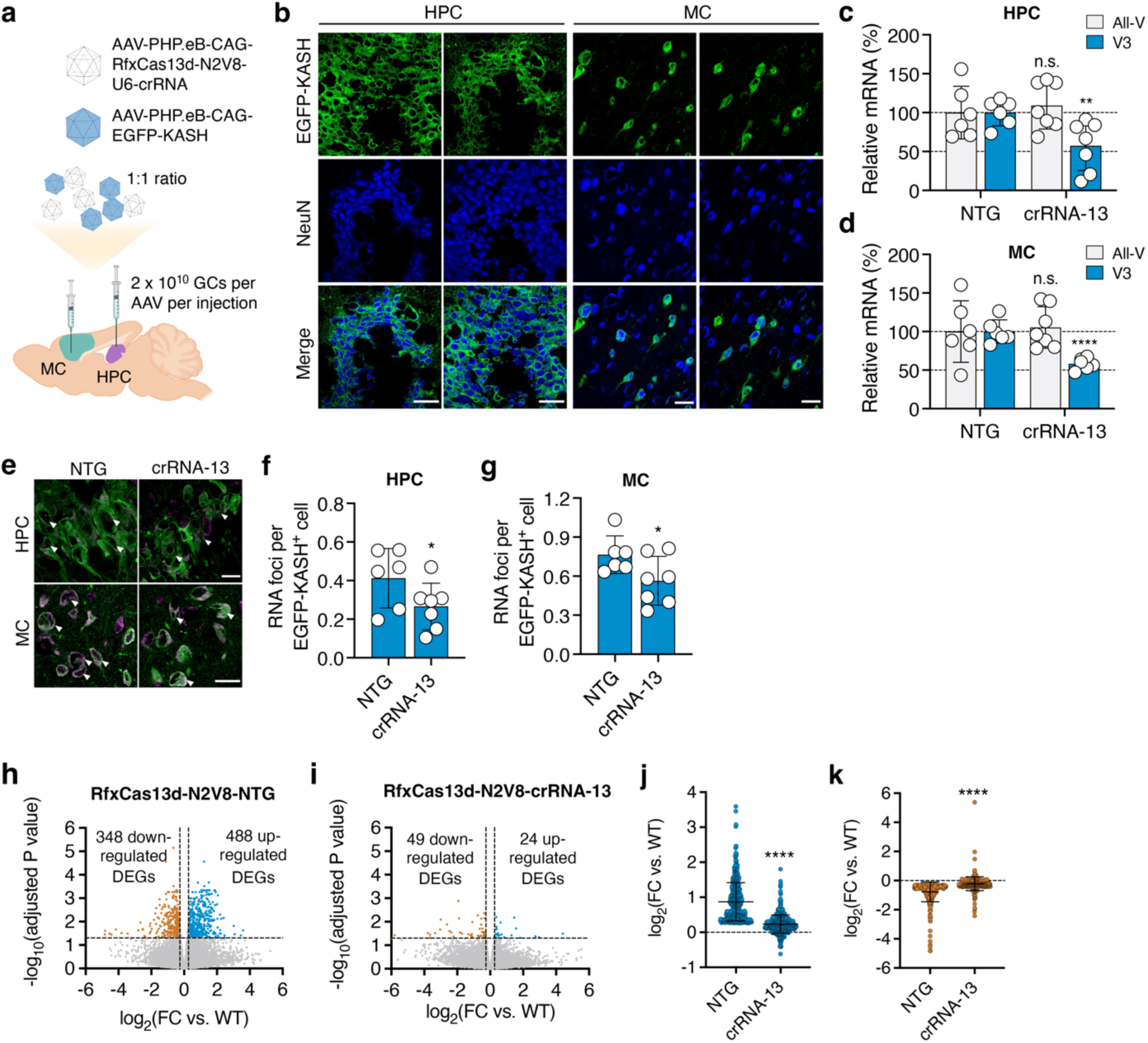
High-fidelity RfxCas13d can target the G_4_C_2_ repeat-containing RNA and reverse deficits in C9-BACexp mice. **(a)** Cartoon illustrating the injection scheme. **(b)** Representative immunofluorescent staining of the hippocampus (HPC) and motor cortex (MC) in C9-BACexp mice two-months after injection with 2 x 10^10^ GCs each of AAV-PHP.eB-RfxCas13d-N2V8-crRNA and AAV-PHP.eB-EGFP-KASH. Scale bar, 30 µm. **(c, d)** Relative all-V and V3 mRNA in EGFP-KASH^+^ nuclei from **(c)** the HPC and **(d)** MC of C9-BACexp injected with AAV-PHP.eB-RfxCas13d-N2V8-crRNA-13 or -NTG with AAV-PHP.eB-EGFP-KASH (*n* ≥ 6). **(e)** Representative FISH for the G_4_C_2_ repeat RNA (purple) in EGFP-KASH^+^ cells from the HPC and MC of C9-BACexp mice injected with AAV-PHP.eB-RfxCas13d-N2V8-crRNA-13 or -NTG and AAV-PHP.eB-EGFP-KASH. Arrowheads indicate representative cells. Scale bar, 20 µm. **(f, g)** Quantification of the number of RNA foci per EGFP-KASH^+^ cell in the **(f)** HPC and **(g)** MC of injected C9-BACexp mice (*n* ≥ 6). **(e-g)** 83-334 and 108-237 cells were counted per animal for the HPC and MC, respectively. 1,432 and 1,016 cells were counted for AAV-PHP.eB-RfxCas13d-N2V8-crRNA-13 and -NTG, respectively, for the HPC, and 1,139 and 919 cells were counted for AAV-PHP.eB-RfxCas13d-N2V8-crRNA-13 and -NTG, respectively, for the MC (*n* ≥ 6). **(h, i)** Volcano plot of the RNA-seq analysis from C9-BACexp mice injected with **(h)** AAV-PHP.eB-RfxCas13d-N2V8-NTG or **(i)** -crRNA-13 with AAV-PHP.eB-EGFP-KASH and compared to wild-type littermates (*n* = 3-7). Lines denote a >1.2-fold change (FC) and an FDR-adjusted P < 0.05. **(j, k)** FC of the **(j)** up-regulated or **(k)** down-regulated DEGs from the analysis in **(h)**. Values indicate means and error bars indicate SD. *P < 0.05, **P < 0.01, ****P < 0.0001; **(c, d, f, g)** one-tailed unpaired t-test; **(j, k)** two-tailed unpaired t-test. **(c, d, f, g)**. All tests compared crRNA-13 to the NTG crRNA. All data points are biologically independent samples.

Consistent with our earlier study, we observed robust EGFP-KASH expression in C9-BACexp mice, finding at two-months post-injection that ∼87% and ∼49% of the cells positive for NeuN in the HPC and MC, respectively, were positive for EGFP-KASH (**Fig. 4b** and **Supplementary Fig. 7**) and that ∼81% and ∼78% of the EGFP-KASH^+^ cells in the HPC and MC, respectively, were positive for RfxCas13d-N2V8 via its HA epitope (**Supplementary Fig. 8**). As before, we observed limited delivery to glial cells, which included GFAP^+^ astrocytes and Iba1^+^ microglia (**Supplementary Fig. 9)**.

Following their isolation by FACS, we used qPCR to measure targeting in EGFP-KASH^+^ nuclei. Compared to control cells, C9-BACexp mice injected with AAV-PHP.eB-RfxCas13d-N2V8-crRNA-13 had a two-fold decrease in V3 mRNA, both in the HPC and the MC (P < 0.01 for the HPC and P < 0.0001 for the MC; **Fig. 4c-d**) and showed no difference in all-V in either region (P > 0.1 for both the HPC and MC; **Fig. 4c-d**), indicating that RfxCas13d-N2V8 preferentially targeted V3 *in vivo*.

Using FISH, we next determined if RfxCas13d-N2V8 affected RNA foci formation in C9-BACexp mice. Within the HPC, we measured a ∼37% decrease in RNA foci positive for the G_4_C_2_ repeat RNA in EGFP-KASH^+^ cells (P < 0.05; **Fig. 4e-f**), with a shift in the distribution of the number of foci per cell for the C9-BACexp mice treated with RfxCas13d-N2V8 (**Supplementary Fig. 10a**). Further, we quantified RNA foci in the MC, finding that C9-BACexp mice treated with RfxCas13d-N2V8 had a ∼27% decrease in the average number of foci per cell (P < 0.05; **Fig. 4e, g**) alongside a shift in their distribution (**Supplementary Fig. 10b)**.

Finally, we evaluated if RfxCas13d-N2V8 could reverse the transcriptional abnormalities that manifest in C9-BACexp mice^34^. To determine this, we injected the MC of six-week-old C9-BACexp mice and their wild-type C57BL/6J littermates with 2 x 10^10^ GCs of AAV-PHP.eB-RfxCas13d-N2V8-crRNA-13 or AAV-PHP.eB-RfxCas13d-N2V8-NTG alongside 2 x 10^10^ GCs of AAV-PHP.eB-EGFP-KASH, which we used to facilitate the isolation of transduced nuclei by FACS at two-months post-injection. We then conducted RNA-seq on the EGFP-KASH^+^ nuclei isolated from each animal.

To uncover the gene alterations that could be attributed to the repeat expansion, a pairwise DEG analysis was conducted between C9-BACexp mice and their C57BL/6J littermates, which were injected with the same vector formulation. In total, we identified 836 genes whose expression was altered in C9-BACexp mice (>1.2-FC; FDR-adjusted P < 0.05; **Fig. 4h** and **Supplementary Data 6 and 7**), 488 of which were up-regulated (**Fig. 4h**), with an average FC from wild-type of 1.83 (**Fig. 4j**) and 348 of which were down-regulated (**Fig. 4h**), with an average FC from wild-type of 1.71 (**Fig. 4k**). Among the DEGs, enrichment (P < 0.05) was observed for terms related to autophagy, RNA splicing and RNA regulation (**Supplementary Data 8**), which have each been linked to C9ORF72 and/or the repeat expansion.

To determine whether RfxCas13d-N2V8 targeting had an effect on the C9-ALS/FTD-associated DEGs, we next conducted a pairwise analysis between C9-BACexp and C57BL/6J mice co-injected with AAV-PHP.eB-RfxCas13d-N2V8-crRNA-13 and AAV-PHP.eB-EGFP-KASH. Of the 836 genes whose expression was altered in C9-BACexp mice injected with the non-targeted crRNA, we found that only 23 of them (∼3%) were affected (that is, deviated significantly from wild-type; >1.2-FC; FDR-adjusted P < 0.05) in C9-BACexp mice treated with crRNA-13 (**Fig. 4i** **and Supplementary Data 6 and 7**), with the 488 and 348 DEGs originally found to be up-and down-regulated measured to deviate from wild-type by a FC of only 1.18 and 1.16, respectively, in C9-BACexp mice treated with crRNA-13 (P < 0.0001 for both compared to NTG; **Fig. 4j-k**). Notably, of these 488 and 348 up-and down-regulated DEGs, we found that 401 (82%) and 253 (73%) of them, respectively, reverted back to wild-type by a FC difference of at least 50%, and that 247 (51%) and 144 (41%) of them, respectively, reverted back to wild-type by a FC difference of at least 75% (**Fig. 4j-k** **and Supplementary Data 7**). These results thus demonstrate that RfxCas13d can at least partially reverse a percentage of the transcriptional deficits that manifested in C9-BACexp mice.

In conclusion, we find that RfxCas13d and a high-fidelity version of it can be used to curb the production of the G_4_C_2_ repeat-containing RNA, while preserving normal C9ORF72 mRNA levels. We further show that RfxCas13d targeting can improve abnormalities associated with the repeat expansion in C9-BACexp mice. These results altogether illustrate the potential of CRISPR-Cas13 technology for C9ORF72-linked ALS/FTD.

## DISCUSSION

An abnormal expansion of a G_4_C_2_ repeat in the first intron of the C9ORF72 gene is the most common genetic cause of ALS^7,8^, accounting for up to 40% of familial forms of the disease and 5-10% of all sporadic cases of the disorder in the United States, Europe and Australia^61^. To date, three non-exclusive mechanisms have been proposed to explain the pathogenicity of the repeat expansion^9–24^. These include haploinsufficiency or loss of C9ORF72 protein functon^9^ and/or acquired toxicity from the effects of bidirectionally transcribed repeat-containing RNAs^13,15–18^ and/or their non-canonical translation to one of five DPR proteins^19–22^.

Given the role that the repeat RNAs may play in C9-ALS/FTD, gene silencing has emerged as a promising strategy for the disorder. Accordingly, both ASOs^27,31,32,59,62^ and miRNAs^25,26^ have both been used to silence the hexanucleotide repeat-containing RNA. These strategies, however, possess limitations that could limit their effectiveness for C9-ALS/FTD. For example, ASOs have a transient lifecycle, which can require a lifetime of administrations to sustain a therapeutic effect^63,64^. This can lead to periods of diminished activity and impose a physical burden on patients. Conversely, while miRNAs can be expressed for an extended period of time from a viral vector, they rely on endogenous RNA processing pathways that are predominately located in the cytoplasm^65–67^ which, in the case of C9-ALS/FTD, could limit their effectiveness, as the repeat RNAs and the RNA foci are located predominately in the nucleus^27,68,69^

One emerging technology whose properties could overcome these limitations is CRISPR-Cas13, particularly RfxCas13d, a class II, type VI CRISPR effector protein that is: (i) capable of cleaving RNAs via its intrinsic RNase activity^33^, (ii) is compact enough to be encoded within a single AAV vector^33^ for a potential single-dose treatment and (iii) can be modified to access the nucleus^33,45^ to engage with the repeat-containing RNAs. Given these properties, we hypothesized that RfxCas13d could be used to target the G_4_C_2_ repeat-containing RNA to influence pathological hallmarks associated with C9-ALS/FTD.

In the present study, we establish a CRISPR-Cas13-based system for C9-ALS/FTD. Using a dual-luciferase reporter screen designed to assess target engagement and collateral effects, we identify crRNAs for RfxCas13d that facilitate the efficient targeting of the G_4_C_2_ repeat-containing RNA in HEK293T cells, SH-SY5Y cells and motor neuron-like cells derived from a patient with C9-ALS. We also find that RfxCas13d effectively curbed the expression of the repeat-containing RNA in C9-BACexp mice, which harbor-the full-length human C9ORF72 gene with ∼100-1,000 copies of the G_4_C_2_ repeat. Our results thus illustrate the potential of CRISPR-Cas13 technology for C9-ALS/FTD.

Critically, after engaging with its target transcript, RfxCas13d undergoes a conformational change that appears to unlock an intrinsic ability to *trans*-cleave non-target RNAs^47,48,70^, a deleterious trait that can limit its applications. As prior studies have demonstrated this property can, in some cases, depend on the crRNA^49,50^, we hypothesized that implementing a screen with a dedicated readout for *trans*-cleavage could aid in the identification of crRNAs that minimize this effect. To this end, we developed a dual-reporter system for identifying effective C9ORF72-targeting crRNAs. This platform consisted of: (i) a Renilla luciferase transgene whose 3’ UTR was fused to the target region of the C9ORF72 gene, and (ii) a constitutively expressed firefly luciferase transgene, whose expression served as a proxy for collateral *trans*-cleavage. As hypothesized, this platform enabled the identification of crRNAs that could effectively target the Renilla luciferase mRNA without appreciably decreasing firefly luciferase, indicating its utility for identifying crRNAs that exert relatively minimal collateral effects in cells.

Given concerns for collateral effects, we also incorporated into our platform engineered variants of RfxCas13d with reduced *trans*-cleavage capabilities, including RfxCas13d-N2V8^50^. According to RNA-seq, RfxCas13d-N2V8 affected the expression of a relatively limited number of genes when programmed to target C9ORF72 (230 DEGs for RfxCas13d-N2V8 versus 1214 DEGs for RfxCas13d). Further, through an over-representation analysis of GO and BP terms, no significant enrichment for any biological themes (FDR-adjusted P > 0.05) was observed among the DEGs affected by RfxCas13d-N2V8, whereas ten functions were enriched (FDR-adjusted P < 0.05) in cells transfected with RfxCas13d. Thus, coupling a dual-reporter system with high-fidelity forms of Cas13 can be an effective means for establishing CRISPR-Cas13-based platforms.

Using FISH, we determined that both RfxCas13d and a high-fidelity counterpart reduced the formation of RNA foci positive for the G_4_C_2_ repeat RNA in C9-BACexp mice, which indicates the ability of these variants to influence a hallmark of C9-ALS/FTD. However, we did not determine if their targeting activity also reduced a DPR protein which, due in large part to their limited abundance, are most reliably quantified using specialized immunoassay platforms^14,32,71,72^. Instead, we show by RNA-seq that RfxCas13d-N2V8 could revert a majority of the transcriptional abnormalities found to manifest in C9-BACexp mice, mirroring a similar finding from a recent study that used RfxCas13d to target huntingtin^38^. This finding is also consistent with a study that previously demonstrated that an ASO for the G_4_C_2_ repeat-containing RNA could at least partially reverse a large percentage of the transcriptional deficits found to manifest in iPSC-derived motor neuron-like cells from an C9-ALS patient^59^.

For this proof-of-concept, we used an AAV vector variant to deliver RfxCas13d to the HPC and MC of C9-BACexp mice, where RfxCas13d was expressed primarily in neurons. While neurons are principally affected in C9-ALS/FTD, other cell types, including astrocytes^73^ and microglia^74^, have also been implicated in the disorder. Thus, additional studies are needed to identify the ideal AAV capsid, and delivery route, for maximizing transduction to all relevant cell populations for C9-ALS/FTD. In addition, we used RfxCas13d to target the sense (GGGGCC) transcript but not the antisense (CCCCGG) variant, which is also thought to contribute to disease pathogenesis^15–18^. Because RfxCas13d is a multiplexable enzyme^33^, it should be feasible to simultaneously target both transcripts in the future. This study is therefore a critical step in a process that we expect to culminate in the development of a dual-targeting platform(s) that rely on high-fidelity forms of Cas13 or alternate RNA-targeting CRISPR effector proteins^75^ to perturb in parallel the sense and antisense repeat-containing RNAs.

In conclusion, we establish a high-fidelity CRISPR-Cas13 system to curb the expression of the G_4_C_2_ repeat-containing RNA for C9-ALS/FTD. Our results illustrate the potential of RNA-targeting CRISPR technologies for C9-ALS/FTD and reinforce the applicability of CRISPR-based approaches for ALS^37,40,76–79^.

## METHODS

### Plasmid construction

The plasmid pAAV-CBh-RfxCas13d-U6-crRNA vector was constructed as previously described^37^.

To construct pSV40-RLuc, a 27-nucleotide spacer sequence was inserted between the BbsI and ClaI restriction sites of psiCHECK-2 (Promega) to remove the firefly luciferase gene sequence and its promoter sequence. A fragment of the C9ORF72 gene encoding 20 copies of the G_4_C_2_ hexanucleotide repeat with 250-and 98-bps of the flanking upstream and downstream sequences from the C9ORF72 gene, respectively, was synthesized (Genscript) and then inserted between the XhoI and NotI restriction sites of the modified psiCHECK-2 plasmid.

To construct pHSV-TK-FLuc, a 32-nucleotide spacer sequence was inserted between the BglII and BbsI restriction sites of psiCHECK-2 to remove Renilla luciferase and its promoter sequence.

To construct pAAV-CBh-RfxCas13d-N2V7-U6-cRNA and pAAV-CBh-RfxCas13d-N2V8-U6-crRNA, the native RfxCas13d gene sequence was PCR amplified from pAAV-CBh-RfxCas13d-U6-crRNA as two fragments using the primers: (1) Fusion-NcoI-Fwd-v2 and Fusion-N2V7-Rev or Fusion-N2V8-Rev; and (2) Fusion-N2V7-Fwd or Fusion-N2V8-Fwd with Fusion-SacI-Reverse (**Supplementary Table 1)**. The resulting fusion PCR product were then ligated into the NcoI and SacI restriction sites of pAAV-CBh-RfxCas13d-U6-crRNA.

crRNAs were cloned as previously described^37^. Briefly, oligonucleotides encoding the crRNA targeting sequences were synthesized (Integrated DNA Technologies) and incubated with T4 polynucleotide kinase (NEB) for 30 min at 37°, incubated at 95°C for 5 min, and then cooled to 4°C at a rate of -0.1°C/s. The duplexed and phosphorylated oligonucleotides were then ligated into the BbsI restriction sites of pAAV-CBh-RfxCas13d-U6-cRNA, _p_AAV-CBh-RfxCas13d-N2V7-U6-cRNA, and pAAV-CBh-RfxCas13d-N2V8-U6-cRNA.

Sanger sequencing (ACGT) was used to confirm the identity of all plasmids. All primer sequences are provided in **Supplementary Table 1.**

### Cell culture, transfections, and luciferase measurements

HEK293T cells and SH-SY5Y cells were cultured in Dulbecco’s modified Eagle’s medium (DMEM; Corning) supplemented with 10% (v/v) fetal bovine serum (FBS; Gibco) and 1% (v/v) antibiotic-antimycotic (Gibco) in a humidified 5% CO_2_ incubator at 37°C.

For the dual-luciferase screen, HEK293T cells were seeded onto a 96 well plate at a density of 2 × 10^4^ cells per well and transfected the following day with 100 ng of pAAV-CAG-RfxCas13d-U6-crRNA, 1 ng of pSV40-RLuc and 10 ng of pHSV-TK-FLuc using polyethylenimine (1 mg/mL; PEI).

At 72 hr post-transfection, HEK293T cells were lysed with Passive Lysis Buffer (Promega). Renilla and firefly luciferase luminescence was then measured using the Dual-Glo Luciferase Assay System (Promega) with a Synergy HTX Plate Reader (BioTek). Renilla and firefly luminescence values for each sample were normalized to values from cells transfected with RfxCas13d with a non-targeted crRNA.

For targeting V3 and all-V, HEK293T cells and SH-SY5Y cells were seeded onto a 24-well plate at a density of 2 × 10^5^ cells per well and transfected the following day with 1 μg of pAAV-CAG-RfxCas13d-U6-crRNA, pAAV-CAG-RfxCas13d-N2V8-U6-crRNA, or pAAV-CAG-RfxCas13d-N2V7-U6-crRNA using Lipofectamine 3000 (ThermoFisher Scientific), according to the manufacturer’s instructions.

### qPCR

RNA from cells or tissue was purified using the RNeasy Plus Mini Kit (Qiagen) and immediately converted to complementary DNA (cDNA) using an iScript cDNA Synthesis Kit (BioRad). qPCR was conducted on a 96-well plate using 30 ng of cDNA with TaqMan Fast Advanced Master Mix (ThermoFisher Scientific) with probes for human HPRT1 (Hs02800695_m1; ThermoFisher Scientific), mouse HPRT1 (MM01318743_M1; ThermoFisher Scientific), C9ORF72-all-V (Hs00376619_m1; ThermoFisher Scientific) and C9ORF72-V3 (Hs00948764_m1; ThermoFisher Scientific). Reaction volumes were 20 µL and probe concentrations were 150 nM and 250 nM for HPRT1 and C9ORF72-all-V and -V3, respectively. C9ORF72 measurements were normalized to HPRT1 for each respective sample. All qPCR reactions were conducted as recommended by the manufacturer’s instructions (ThermoFisher Scientific). CBLN1 and CBLN2 measurements were performed using iTaq Universal SYBR Green Supermix (BioRad) and normalized to human GAPDH.

### Western blot

Cells were lysed by radioimmunoprecipitation assay (RIPA) buffer [0.2% IGEPAL CA-620, 0.02% SDS with Protease Inhibitor Cocktails (VWR Life Science, 97063-010)]. Protein concentration was then determined using the DC Protein Assay Kit (Bio-Rad). A total of 20 μg of protein per sample was electrophoresed by SDS-PAGE and electrophoretically transferred to a polyvinylidene fluoride (PVDF) membrane in transfer buffer [20 mM Tris-HCl, 150 mM glycine, and 20% (v/v) methanol] for 30 min at 100 V using a Criterion Blotter (Bio-Rad). Membranes were blocked with 5% (v/v) blotting-grade blocker (Bio-Rad) in TBS (10 mM Tris-HCl and 150 mM NaCl, pH 7.5) with 0.05% Tween-20 (TBS-T) for 1 hr and then incubated with primary antibody in blocking solution at 4 °C overnight. The following primary antibodies were used: Rabbit anti–β-actin (1:1000; Cell Signaling Technology, 4970S) and rabbit anti-C9ORF72 (1:1000; Proteintech, 22637-1-AP).

After incubation with primary antibody, membranes were washed three times with TBS-T and incubated with goat anti-rabbit horseradish peroxidase conjugate (1:4000; ThermoFisher Scientific, 65-6120) in blocking solution for 1 hr at room temperature (RT). Membranes were then washed three final times with TBS-T and treated with SuperSignal West Dura Extended Duration Substrate (Thermo Fisher Scientific). Chemiluminescence was detected using a ChemiDoc XRS+ (Bio-Rad). Band intensities were quantified using Image Lab Software (Bio-Rad) and normalized to the reference protein in each sample.

### RNA sequencing

Library construction was conducted by the Roy J. Carver Biotechnology Center (University of Illinois Urbana-Champaign, Urbana, IL) as previously described^37^. Briefly, RNAs were purified using the RNeasy Plus Mini Kit (Qiagen) and subsequently treated with DNase. RNAs were then converted into individually barcoded polyadenylated mRNA sequencing libraries using the Kapa Hyper Stranded mRNA library kit (Roche) and fused with unique dual indexes. Adaptor-ligated double-stranded cDNAs that were synthesized from the mRNA libraries were then PCR-amplified for eight cycles with KAPA HiFi DNA Polymerase (Roche), quantitated by PCR and subsequently pooled in equimolar concentration. The libraries were then sequenced by a NovaSeq 6000 (Illumina) using 2x150nt reads on an S1 lane. The FASTQ files that were generated from the sequencing were demultiplexed using the bcl2fastq v2.20 Conversion Software (Illumina). The quality of the demultiplexed FastQ files was then evaluated using FastQC (version 0.11.9).

Mouse RNAs were purified using the Rneasy Plus Mini Kit (Qiagen) and subsequently treated with DNase. Libraries were then prepared using the Universal RNA-Seq Kit (Tecan) with probes to deplete mouse rRNAs, with the quality of each sample assessed using a 5200 Fragment analyzer (Agilent). The final barcoded RNA-seq libraries were pooled in equimolar concentration and sequenced by a NovaSeq 6000 (Illumina) using 1x100nt reads on one S2 lane for 101 cycles. The FASTQ files that were generated from the sequencing were demultiplexed using the bcl2fastq v2.20 Conversion Software (Illumina). The quality of the demultiplexed FastQ files was then evaluated using FastQC (version 0.11.9).

Data analysis was conducted by the High-Performance Biological Computing Core (University of Illinois Urbana-Champaign, Urbana, IL). Salmon (version 1.5.2 for the human study and version 1.10.0 for the mouse study) was used to quasi-map reads to the transcriptome and to quantify the abundance of each transcript. For the human study, spliced transcript sequences from Annotation Release 109.2020112 (NCBI) were used along with the GRCh38 reference genome as the decoy sequence for the Salmon index. Because reads from the mouse study contained a higher proportion retained introns, both spliced and un-spliced transcripts from Annotation Release 109 (NCBI) was used with the three main human C9ORF72 transcript variants and EGFP-KASH with the GRCm39 reference genome as the decoy sequence for the Salmon index. Quasi-mapping was then performed in an identical manner for the human and mouse studies with the additional arguments: --seqBias and –gcBias, which were used to correct for sequence-and GC-specific biases, --numBootstraps=30, which was used to compute bootstrap transcript abundance estimates, and –validateMappings and –recoverOrphans, which were used to improve the accuracy of mappings.

Gene-level counts were estimated on transcript-level counts using the “lengthScaled TPM” method from the tximport package^80^ to provide accurate gene-level counts estimates and to keep multi-mapped reads in the analysis. DEG analysis was performed using the limma-trend method plus 1 (human) or 4 (mouse) extra factors estimated by the RUVseq package to correct for spurious technical variation^81^. An FDR adjustment was then conducted globally across the pairwise comparisons. Processed RNA-seq data for the human and the mouse studies are available in **Supplementary Data** 1 and **Supplementary Data 6**, respectively.

Overrepresentation analyses on DEGs was performed using NetworkAnalyst 3.0^82^ and Enrichr^83^ to identify the enriched biological terms associated with the differentially expressed genes in HEK293T and C9-BACexp mouse samples, respectively, using the GO:BP databases^84^.

### AAV packaging

AAV vectors were packaged as described^85^. Briefly, 2 x 10^7^ HEK293T cells were seeded onto 15-cm plates in DMEM supplemented with 10% (v/v) FBS and 1% (v/v) antibiotic-antimycotic (ThermoFisher Scientific). At 16 hr after seeding, cells were transfected with 15 μg of pAAV-CBh-RfxCas13d-U6-crRNA-13, -7, -1 and -NTG, pAAV-CBh-RfxCas13d-N2V8-U6-crRNA-13 and -NTG, or pAAV-CBh-EGFP-KASH with 15 μg of pAAV-PHP.eB and 15 μg of pHelper using 135 μL of polyethylamine (1 μg/ μL).

Cells were harvested by a cell scraper at five-days post-transfection and centrifuged at 4000g for 5 min at RT. Cells were then resuspended in lysis buffer (50 mM Tris-HCl and 150 mM NaCl, pH 8.0) and subsequently freeze-thawed three consecutive times using liquid nitrogen and a 37°C water bath. Afterwards, cells were incubated with benzonase (10 units per 1 mL of suspension; Sigma-Aldrich) for 30 min at a 37°C. The suspension was then centrifuged for 30 min at 18,500g at RT, with the ensuing supernatant layered on an iodixanol density gradient, as described^85^. The iodixanol density gradient was centrifuged for 2 hr at 140,000g at 18°C and virus was extracted, washed three times with 15 mL of PBS with 0.001% Tween-20 and concentrated to less than 250 μL using an Ultra-15 Centrifugal Filter Unit (Amicon). Viral titers were determined by qPCR using iTaq Universal SYBR Green Supermix (Bio-Rad).

### Stereotaxic injections

All procedures were approved by the Illinois Institutional Animal Care and Use Committee (IACUC) at the University of Illinois Urbana-Champaign and conducted in accordance with the National Institutes of Health (NIH) Guide for the Care and Use of Laboratory Animals.

Two-month-old C9-BACexp mice [C57BL/6J-Tg(C9ORF72_i3)112Lutzy/J; Jackson Laboratory, Stock #023099] were injected at stereotaxic coordinates anterior-posterior (AP) = 1.2 mm; medial-lateral (ML) = ±1.4 mm; and dorsal-ventral (DV) = 2 mm and 1.7 mm for the MC and stereotaxic coordinates AP = -1.9 mm; ML = ±1.3 mm; and DV = 1.8 mm and 1.3 mm for the HPC. 1 x 10^10^ GCs of each vector was delivered per depth in 2.4 uL of saline solution for a total of 2 x 10^10^ GCs per animal. Injections were performed using a drill and microinjection robot (NeuroStar).

### Nuclei isolation

Neuronal nuclei were isolated from dissected brain as described^86^. Briefly, tissue from the MC and HPC were dissected and homogenized in 2 mL of Nuclei EZ Lysis Buffer (Sigma-Aldrich) using a KIMBLE Dounce Tissue Grinder (Sigma-Aldrich). After the addition of an additional 2 mL of Nuclei EZ Lysis Buffer, samples were incubated at RT for 5 min. Homogenized tissues were then centrifuged at 500g for 5 min. After removing the supernatant, nuclei were resuspended in 4 mL of Nuclei Suspension Buffer (PBS with 100 µg/mL of BSA) and centrifuged at 500g for an additional 5 min. The resulting pellet was then resuspended in 1 mL of Nuclei Suspension Buffer for FACS. Nuclei were strained through Round-Bottom Polystyrene Test Tubes with a 35 µm Cell Strainer Snap Cap (Falcon) and subjected to FACS using a BD FACSAria II Cell Sorter (Roy J. Carver Biotechnology Center Flow Cytometry Facility, University of Illinois Urbana-Champaign, Urbana, IL). Nuclei were collected in 350 µL of RNeasy Plus Kit Lysis Buffer (Qiagen) and at least 15,000 nuclei were collected for each sample.

### Immunofluorescent analysis

Immunofluorescent analyses were performed as described^76^. Briefly, after their extraction, brains were fixed in 4% paraformaldehyde (PFA) overnight at 4°C. Fixed tissues were then sliced to 40 µm sagittal sections on a CM3050 S cryostat (Leica) and stored in cryoprotectant solution at -20°C. For staining, sections were washed three times in PBS for 15 min and incubated in blocking solution [PBS with 10% (v/v) donkey serum (Abcam) and 0.5% Triton X-100] for 2 hr at RT. Sections were then stained with primary antibodies in blocking solution for 72 hr at 4°C. After incubation, sections were washed three times with PBS and incubated for 2 hr with the secondary antibodies at RT. Sections were then washed three final times with PBS and mounted onto slides using VECTASHIELD HardSet Antifade Mounting Medium (Vector Laboratories).

Sections were imaged using a Leica TCS SP8 confocal microscope (Beckman Institute Imaging Technology Microscopy Suit, University of Illinois Urbana-Champaign, Urbana, IL). Images were analyzed using ImageJ imaging software by a blinded investigator.

Primary antibodies were rabbit anti-HA (1:500; Cell Signaling Technology, 3724S), mouse anti-NeuN (1:1000; Millipore Sigma, MAB377), rabbit anti-Iba1 (1:500; Wako Pure Chemicals Industries, 019-19741), and chicken anti-GFAP (1:1000; Abcam, ab4674).

Secondary antibodies were donkey anti-mouse Alexa Fluor 647 (1:150; Jackson ImmunoResearch, 715-605-151), donkey anti-rabbit Cy3 (1:150; Jackson ImmunoResearch, 711-165-152), donkey anti-rabbit Alexa Fluor 647 (1:150; Jackson ImmunoResearch , 711-605-152), donkey anti-chicken Alexa Fluor 647 (1:150; Jackson ImmunoResearch , 703-605-155) .

## FISH

Tissue sections were incubated in blocking solution [DEPC PBS with 10% (v/v) donkey serum and 0.5% (v/v) Triton X-10] for 2 hr at RT. Sections were then washed three times with DEPC PBS and incubated in hybridization solution [2x DEPC saline-sodium citrate (SSC) with 50% (v/v) formamide, 10% (w/v) dextran sulfate, 50 mM sodium phosphate and 0.5% (v/v) Triton X-100] with 40 nM of the FISH probe for 24 hr in the dark at 37°C. Afterwards, sections were incubated in the dark at 66°C for 2 hr. Tissue sections were then washed once in 2x DEPC SSC and subsequently washed twice with 0.1x DEPC SSC at RT. Tissues were stained with DAPI, mounted onto slides with VECTASHIELD HardSet Antifade Mounting Medium (Vector Laboratories) and stored at 37°C. Sections were then imaged using a Leica TCS SP8 microscope (Beckman Institute Imaging Technology Microscopy Suit, University of Illinois Urbana-Champaign, Urbana, IL). 58-455 EGFP-KASH^+^ cells were counted for each injection site. Images were analyzed using ImageJ by a blinded investigator.

The sequence of the FISH probe used to detect the G_4_C_2_-containing RNA foci was: 5TYE563/CCCCGGCCCCGGCCCC/3TYE563. The probe was previously validated^27^ and custom-synthesized by Qiagen.

### Differentiations

Neural progenitor cells (NPCs) derived from a 74-year-old female ALS patient with >145 copies of the hexanucleotide repeat expansion in the C9ORF72 gene were obtained from AXOL Bioscience (ax0073). Upon arrival, NPCs were immediately resuspended at a concentration of ∼2.6 × 10^5^ cells per mL in Motor Neuron Maintenance Medium (AXOL Bioscience, ax0072) with 0.2 µM Compound E (Abcam, ab142164), 0.1 µM retinoic acid (Sigma-Aldrich, R2625) and 10 µM ROCK Inhibitor (Focus Biomolecules, 10-2301). The cells were then seeded on 24-well plate that was pre-coated with 0.5 mg/mL poly-D-Lysine (Sigma-Aldrich, P7405) and vitronectin (ThermoFisher, A14700) at a density of ∼1.3 × 10^5^ cells per well.

For the differentiation, as specified by the manufactures instructions, a full volume medium exchange was conducted each day using Complete Motor Neuron Maintenance Medium with 0.2 µM Compound E (Abcam, ab142164), 0.5 µM retinoic acid (Sigma-Aldrich, R2625), 10 ng/mL recombinant human ciliary-derived neurotrophic factor (CNTF) (ax139888), 5 ng/mL recombinant human brain-derived neurotrophic factor (BDNF) (ax139800), 10 ng/mL recombinant human brain-derived neurotrophic factor (GDNF) (ax139855), and motor neuron maturation accelerator supplement (ax0179).

At day 14 post-seeding, cells were treated with AAV-PHP.eB vector encoding: (i) RfxCas13d-N2V8 with either crRNA-13 or a non-targeted crRNA or (ii) EGFP-KASH, both in the presence of 1% (v/v) antibiotic-antimycotic (Gibco). Vector was added to cells at an MOI of ∼2 × 10^6^.

### Immunocytochemistry

Differentiated neuron-like cells were fixed in 200 µL of 4% (v/v) PFA for 15 min at RT and subsequently incubated with PBS with 0.1% Triton X-100 for 10 min at RT, after which they were washed three times with PBS. To block, cells were incubated in blocking solution [PBS with 10% (v/v) donkey serum (Abcam) and 0.1% Tween 20] for 30 min at RT. The cells were then incubated with primary antibodies in blocking solution overnight at 4°C. After incubation, cells were washed three times with PBS and incubated with secondary antibodies in blocking solution for 1 hr at RT, after which they were washed three times with PBS.

Primary antibodies were mouse anti-HB9 (1:200; Thermo Fisher Scientific, PA5-23407), and goat anti-ChAT (1:50; EMD Millipore, AB144P.

Secondary antibodies were donkey anti-goat Alexa Fluor 647 (1:150; Jackson ImmunoResearch, 705-605-147), and donkey anti-mouse Cy3 (1:150; Jackson ImmunoResearch, 715-165-150).

Cells were imaged using a Zeiss Observer Z1 microscope (Beckman Institute Imaging Technology Microscopy Suit, University of Illinois Urbana-Champaign, Urbana, IL). Images were processed using ImageJ imaging software.

### Statistical analysis

Statistical analysis was performed using GraphPad Prism 8. Unless otherwise noted, all measurements were compared using a one-tailed unpaired t-test.

## ACKNOWLEDGMENTS

We thank A. Hernandez for helpful discussion and assistance with library preparations, J. Drnevich for assistance with the RNA-seq analyses, and P. Perez-Pinera for access to the Synergy HTX Plate Reader This work was supported by the NIH/NINDS (1U01NS122102-01A1, 1R01NS123556-01A1), the NIH/NIGMS (5R01GM141296), the Muscular Dystrophy Association (MDA602798), the Simons Foundation (887187), the Parkinson’s Disease Foundation (PF-IMP-1950), the Judith & Jean Pape Adams Foundation and the ALS Association (20-IIP-516). M.A.Z.C. was supported by the NIH/NIBIB (T32EB019944), the Mavis Future Faculty Fellows Program, and an Aspire Fellowship from the University of Illinois Urbana-Champaign. Cartoons were created with BioRender.

## AUTHOR CONTRIBUTIONS

T.G. conceived of the study; J.E.P, T.X.M, and S.Z., cloned the plasmids; T.X.M conducted the luciferase assays. T.X.M. conducted the qPCR measurements in HEK293T, SH-SY5Y, FACS-isolated nuclei, and neuron-like cells; C.K.W.L. conducted western blots; T.X.M. packaged the AAV vectors; T.X.M. and C.K.W.L. maintained animal; T.X.M. and M.A.Z.C. conducted the stereotaxic injections; T.X.M. harvested tissues and conducted FACS; T.X.M. and C.K.W.L. conducted the immunofluorescent analyses; W.M.T. analyzed RNA FISH experiments; T.X.M. and C.K.W.L. conducted the neuronal differentiations and conducted immunocytochemistry analyses on the cells; T.G., T.X.M., and C.K.W.L. wrote the manuscript with input from all authors.

